# Vulnerabilities of the SARS-CoV-2 virus to proteotoxicity – opportunity for repurposed chemotherapy of COVID-19 infection

**DOI:** 10.1101/2020.04.07.029488

**Authors:** Maryam Al-Motawa, Hafsa Abbas, Patrick Wijten, Alberto de la Fuente, Mingzhan Xue, Naila Rabbani, Paul J. Thornalley

**Author notes:** Correspondence should be addressed to Paul J. Thornalley. Joint corresponding authors.

## Abstract

The global pandemic of COVID-19 disease caused by infection with the SARS-CoV-2 coronavirus, has produced an urgent requirement and search for improved treatments whilst effective vaccines are developed. A strategy for improved drug therapy is to increase levels of endogenous reactive metabolites for selective toxicity to SARS-CoV-2 by preferential damage to the viral proteome. Key reactive metabolites producing major quantitative damage to the proteome in physiological systems are: reactive oxygen species (ROS) and the reactive glycating agent methylglyoxal (MG); cysteine residues and arginine residues are their most susceptible targets, respectively. From sequenced-based prediction of the SARS-CoV-2 proteome, we found 0.8-fold enrichment or depletion of cysteine residues in functional domains of the viral proteome; whereas there was a 4.6-fold enrichment of arginine residues, suggesting SARS-CoV-2 is resistant to oxidative agents and sensitive to MG. For arginine residues of the SARS-CoV-2 predicted to be in functional domains, we examined which are activated towards modification by MG – residues with predicted or expected low pK_a_ by neighbouring group in interactions. We found 25 such arginine residues, including 2 in the spike protein and 10 in the nucleoprotein. These sites were partially conserved in related *coronaviridae*: SARS-COV and MERS. Finally, we identified drugs which increase cellular MG concentration to virucidal levels: antitumor drugs with historical antiviral activity, doxorubicin and paclitaxel. Our findings provide evidence of potential vulnerability of SARS-CoV-2 to inactivation by MG and a scientific rationale for repurposing of doxorubicin and paclitaxel for treatment of COVID-19 disease, providing efficacy and adequate therapeutic index may be established.

## INTRODUCTION

A global pandemic of COVID-19 disease caused by infection with the SARS-CoV-2 coronavirus has developed from January 2020. It has produced a global public health emergency with currently (20^th^ July 2020) over 14 million infections and *ca*. 600,000 deaths, with both rapidly increasing. New treatments are urgently required for COVID-19 disease until effective vaccines are developed. A rapid route to achieve this is repurposing of existing drugs with previously undisclosed activity against coronavirus infection.

As a strategy to identify drugs for repurposing, we sought to explore whether the SARS-CoV-2 may have vulnerabilities in the viral proteome to modification by endogenous reactive metabolites. Pharmacological increase of reactive metabolites will then produce a virucidal effect and therapeutic response for COVID-19 disease. Important reactive metabolites producing major quantitative modification of the proteome in physiological systems are: reactive oxygen species (ROS) and methylglyoxal (MG) (Winterbourn, 2008;Rabbani and Thornalley, 2015). Key to characterizing the vulnerability of the viral proteome to reactive metabolites, ROS and MG, is location of their susceptible amino acid residue targets in functional domains of viral proteins and activation of these residues towards reaction with reactive metabolites. ROS are formed by mitochondria through trace leakage of electron flux in oxidative phosphorylation, by oxidases and other sources. They are metabolized by antioxidant enzymes, superoxide dismutase, catalase, glutathione peroxidase and peroxiredoxins (Murphy et al., 2011). The reactive dicarbonyl metabolite, MG, is formed mainly by trace level degradation of triosephosphate glycolytic intermediates, glyceraldehyde-3-phosphate and dihydroxyacetonephosphate, and is mainly metabolised by glutathione-dependent glyoxalase 1 (Glo1) of the glyoxalase pathway (Rabbani et al., 2016b) (**Figure 1A**). The most susceptible targets in proteins to modification by ROS are cysteine residues which are oxidised to cystine and cysteine sulfenic and sulfonic acids (Winterbourn, 2008). The most susceptible targets in proteins to modification by MG are arginine residues which are glycated to hydroimidazolone MG-H1 with loss of charge, all related electrostatic interactions and, typically, resistance to proteolytic cleavage close to the site of modification (Rabbani et al., 2016b) (**Figure 1B**).

**Figure 1.**
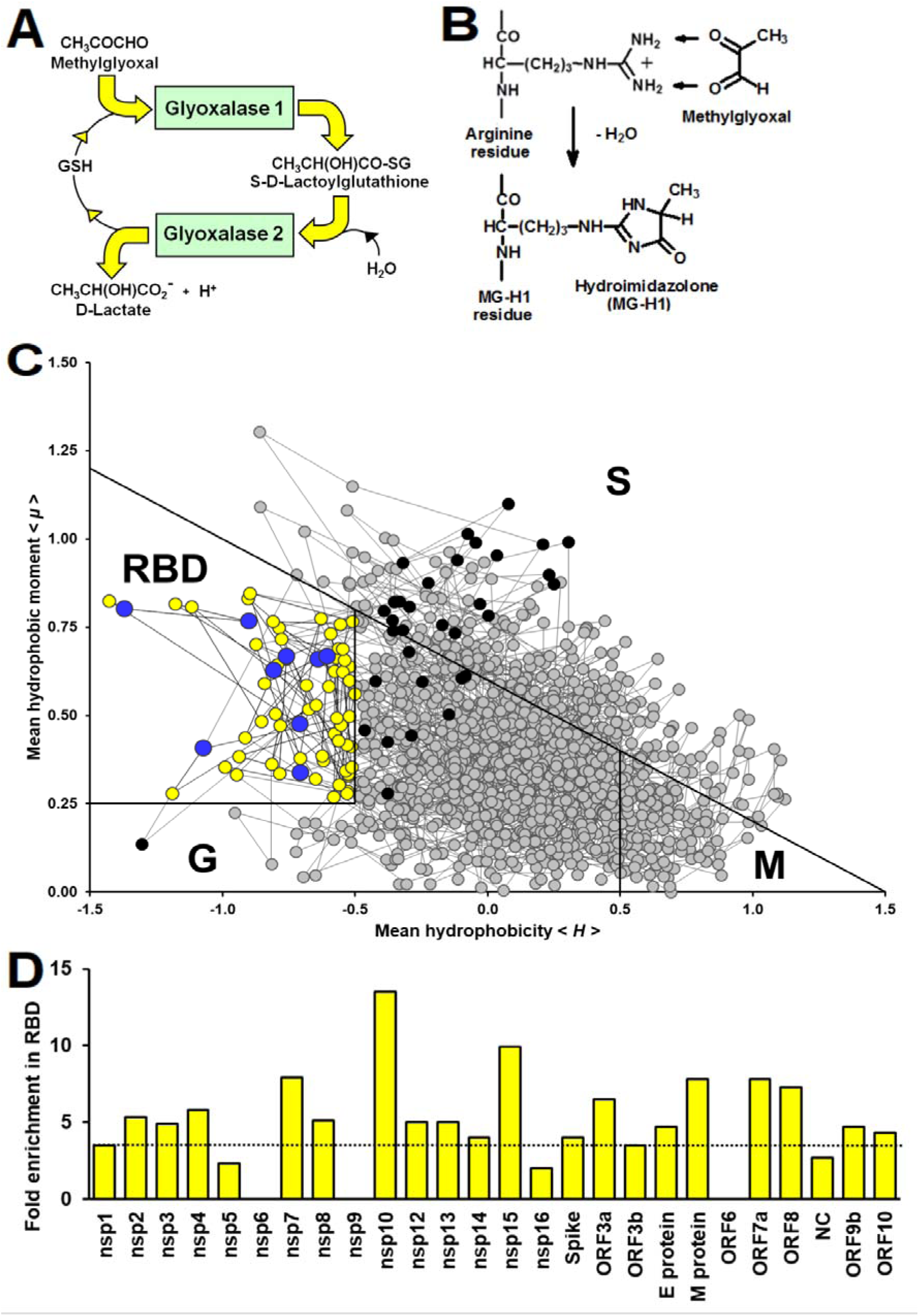
Methylglyoxal – an endogenous arginine-modifying reactive metabolite and receptor binding domain analysis of functional arginines in the SARS-CoV-2 proteome. **(A)** Metabolism of methylglyoxal by the glyoxalase pathway (Rabbani et al., 2016b). (**B)** Modification of arginine residues by MG to form hydroimidazolone, MG-H1. **(C)** Receptor binding domain (RBD) plot for SARS-CoV-2 Spike protein. Line-linked filled circles represent the primary sequence. The RBD is the area bound by the trapezium in the upper left-side region of the chart. Key:◯, arginine residue in the RBD;◯, arginine residues outside the RBD;◯, other amino acid residues in the RBD other amino acid residues in the RBD; and ◯, other amino acid residues outside the RBD. Other predicted domains: surface (S), globular (G) and membrane (M). **(D)** Arginine enrichment in individual proteins of SARS-CoV-2 proteins. Proteins not shown have no arginine residues (nsp11, ORF7b and ORF14).

Key to exploring if reactive metabolites of the host can be exploited to produce a virucidal response against SARS-CoV-2 is to identify proteomic vulnerabilities of the virus. Currently it is unknown if target amino acid residues of reactive metabolites are enriched in functional domains of the viral proteome, and if these targets amino acids are activated towards modification by reactive metabolites. It also unknown if there are investigational new drugs or current clinically approved drugs that increase reactive metabolites to virucidal levels in the cellular environment where SARS-CoV-2 undergo cell fusion and propagation. To address these gaps in knowledge, we initiated a series of studies using bioformatics tools, available proteomics data and a cell model used in SARS-CoV-2 virus propagation. Herein we report the susceptibility of the SARS-CoV-2 virus to increased MG or “dicarbonyl stress” (Rabbani and Thornalley, 2015). This is based on enrichment of arginine residues in functional domains of the SARS-CoV-2 proteome and predicted activation of many of these arginine residues to modification by MG through neighbouring group interactions. We also identified two clinical antitumor drugs that increase the cellular concentration of MG to virucidal levels and are candidates for consideration for repurposing for evaluation for treatment of COVI-19.

## MATERIALS AND METHODS

### Reagents and Chemicals

Doxorubicin, Paclitaxel, monoclonal anti-Glo1 antibody (rat), anti-Rat IgG (whole molecule)–Biotin conjugate, D-Lactic dehydrogenase were purchased from Sigma-Aldrich (Poole, Dorset, UK). Geneticin G-418 (potency rating – 700 µg) was purchased from Fisher Scientific (Loughborough, UK). S-p-bromobenzylglutathione cyclopenyl diester (BBGD) was prepared an purifeid in-house, as described (Thornalley et al., 1996). The HEK293 cell line was purchased from the American Tissue Culture Collection (ATCC, Virginia, USA). Plasmids, pIRES2-GLO1-EGFP and pIRES-EGFP, PJT laboratory were prepared and purified in-house, as described (Ahmed et al., 2008).

### Sequences of SARS CoV-2, SARS-CoV and MERs and human host proteins

Reference sequences of the 29 proteins of the SARS-CoV-2 proteome (Table S1) and sequences of analogous proteins of SARS-CoV and Middle East Respiratory Syndrome (MERS) *coronaviridae* were obtained from the NBCI reference sequence database (www.ncbi.nlm.nih.gov). Sequences of reviewed proteins of the human proteome, 18,821 – excluding fetal proteins, were obtained from the UniProtKB database (www.uniprot.org).

### Receptor binding domain analysis

Receptor binding domain (RBD) analysis is a protein primary sequenced based informatics method to deduce amino acid residues in functional domains of proteins – defined as sites of protein-protein, protein-nucleic acid and protein-ligand or substrate interaction. It is applicable to any protein. The optimised protocol uses a window of 5 amino acid residues moved sequentially along the sequence of a protein, assuming a gyration angle between two consecutive residues in the sequence of 100°, to deduce sequential mean Eisenberg hydrophobicity and mean dipole moment for the central amino acid. Values cannot be deduced for the two amino acids at the N- and C-termini of proteins and they are therefore missing from the amino acid residue prevalence reports (Gallet et al., 2000). This approach had 80% accuracy when validated against a database of known interacting proteins (Gallet et al., 2000). We developed an R script to obtain mean hydrophobicity and hydrophobic moment for all UniProtKB proteins and SARS-CoV-2 proteins.

### Prediction of arginine residues activated for methylglyoxal modification

Arginine residues of proteins are activated towards reaction with MG by decrease of pK_a_ of the guanidino side chain which facilitates formation of the MG-guanidino sidechain encounter (Rabbani and Thornalley, 2012;Rabbani et al., 2016a). Arginine residue sidechain pK_a_ is decreased by interaction with neighbouring amino acid residues with positively charged sidechains. For an α-helix, interactions with lysine or arginine residues at positions - 4, -3, +3 and +4 in the sequence with respect to the target arginine residue are expected to decrease the arginine target residue pK_a_ by side chain interaction along the side of the helix axis. Longer range interactions occur between these and other types of secondary structure domains in the tertiary structures of proteins where arginine residue pK_a_ may be predicted from crystallographic data. We explored the peptide environments of arginine residues in predicted functional domains of the SARS-CoV-2 proteome, identifying arginine residues with neighbouring interacting lysine and arginine residues, predicted secondary structure and predicted arginine residues target pK_a_ where crystallographic data are available. Information om predicted secondary structure was extracted from *in silico* predicted models: nsp1, YxJyvF; nsp3, 5hYU6g; M-protein, 9LzAZz (http://3dbionotes.cnb.csic.es/ws/covid19) (Waterhouse et al., 2018). Arginine side chain predicted pK_a_ values are given, deduced by DelPhiPKa program (Wang et al., 2016) using nucleoprotein crystal structure (pdb file 6VYO; Chang *et al*., to be published) and AMBER forcefield (predictions were similar with CHARMM and PARSE forcefields). To identify similar arginines residues in SARS-CoV and MERS proteins, we used the Clustal Omega software on-line (Madeira et al., 2019).

### Culture of HEK293 cells *in vitro*

The HEK293 cell line, seeding density 2 × 10^4^ cells cm^-2^, was cultured in Dulbecco’s Modified Eagles Medium (DMEM) containing phenol red, L-glutamine and 4500 mg/L glucose, supplemented with 10% Foetal Bovine Serum (FBS), 100 U penicillin and 0.1 mg/ml streptomycin. pIRES2-GLO1-EGFP plasmid (Glo1+ vector) and pIRES-EGFP plasmid (empty vector) were prepared as described (Ahmed et al., 2008). HEK293 cells were stably transfected with Glo1+ and empty vector using Lipofectamine 2000, according to the manufacturer’s instructions (plasmid DNA: Lipofectamine 2000, 1:4). After 48 h, cells were sub-cultured, G-418 disulphate was added (2 mg/ml; 405 µg/mg potency) culture continued. Transfected colonies with GFP fluorescence were selected using a cloning disc (3.2 mm) and glass cylinder selector (8 mm, 150 µl) and cultured further with G-418 disulphate (1 mg/ml, 705 µg/mg potency) containing medium. Assessment of Glo1 activity and protein, as described (Arai et al., 2014;Xue et al., 2014), indicated a 4 - 5 fold increase in Glo1 activity and protein. HEK293 cells stably tranfected with empty and GLO1+ vectors were incubated with and without cell permeable Glo1 inhibitor, S-p-bromobenzylglutathione cyclopenyl diester (BBGD) (Thornalley et al., 1996), doxorubucin and paclitaxel at the concentrations indicated (diluted from 100 mM stock solution in DMSO) for 2 days and effect on cell growth assessed by viable cell number counts, using the Trypan blue exclusion method and median growth inhibitory concentrations GC_50_ deduced. Cellular MG concentration and flux of formation of D-lactate, a surrogate measure of flux of formation of MG, was assayed as described (Rabbani and Thornalley, 2014;Irshad et al., 2019).

### Statistical analysis

Datasets were checked for normality of distribution and parametric statistical tests for assessment of significance of difference between study groups applied: *Student’s t-test* for two groups and one-way *ANOVA* for 3 or more study groups.

## RESULTS

### Enrichment of arginine residues in the functional domains in the SARS-CoV-2 proteome

We acquired primary amino acid sequences of the 29 proteins of the SARS-CoV-2 proteome (**Table S1**) and also, for comparison, 18,821 reviewed protein sequences of human host proteins from the UniProt Knowledgebase (UniProtKB; www.uniprot.org). We found a similar prevalence of cysteine and arginine residues in the viral proteome: 3.14% and 3.63%, respectively (**Table 1**). We applied RBD analysis to identify functional domains of viral proteins and to thereby deduce the prevalence and enrichment of cysteine and arginine residues therein. The RBD analysis outcome is illustrated as a plot of mean hydrophobicity against mean dipole moment for the widow of 5 amino acid residues moved sequentially along the sequence of a protein. An example of the RBD analysis of the SARS-CoV-2 Spike protein is given in **Figure 1C**. Functional domains are located in a trapezium-shaped domain on the top-left side of the plot – regions of low mean hydrophobicity and high mean dipole moment of proximate groups of amino acid residues. This analysis showed that 4.8% of cysteine residues were in functional domains of the SARS-CoV-2 proteome whereas a much greater proportion of arginine residues, 30.7%, were in functional domains. The enrichment of arginine residues in functional domains was 4.9-fold – the highest of any amino acid, whereas there was a slight negative enrichment, 0.8-fold, or depletion of cysteine residues in functional domains. Other amino acid residues susceptible to oxidative damage were also depleted in functional domains: met 0.6, tyr 0.8 and trp 0.3 (**Table 1**). The SARS-CoV-2 proteome is, therefore, resistant to oxidative inactivation but susceptible to functional inactivation by MG. For individual SARS-CoV-2 proteins, the majority had arginine residue enrichment in the functional domains greater than the mean of the human host proteome of *ca*. 3.5: range 2.3 – 13.5 (**Figure 1D**).

**Table 1.** Receptor binding domain of SARS-CoV-2 viral proteomes.

| Amino<br>acid | N |  | Prevalence |  | Proportion | Fold |
| --- | --- | --- | --- | --- | --- | --- |
|  | All | RBD | All | RBD | in RBD (%) | Enrichment |
| Arg | 358 | 110 | 3.63 | 17.7 | 30.7 | 4.9 |
| Cys | 310 | 15 | 3.14 | 2.4 | 4.8 | 0.8 |
| Met | 203 | 7 | 2.06 | 1.1 | 3.5 | 0.6 |
| Tyr | 448 | 23 | 4.54 | 3.7 | 5.1 | 0.8 |
| Trp | 113 | 2 | 1.15 | 0.3 | 1.8 | 0.3 |

### Arginine residues activated for methylglyoxal modification by predicted neighboring group interaction

We next sought to identify of arginine residues in the predicted functional domains of SARS-CoV-2 proteins are activated towards modification by MG based on potentially neighbouring group side-chain interaction with arginine and lysine residues and, where crystallographic data are available, predicted target arginine residue pK_a_. For example, in human serum albumin, neighbouring group interactions with R186, R218 and R410 decrease the pK_a_ values of their sidechain guanidino groups to 12.5, 12.2 and 12.5 from the basal pK_a_ of 13.8 (Fitch et al., 2015). The reactivity with MG of these arginine residues increases by 20 – 40 fold through increase of the trace level conjugate base of the sidechain guanidino group (Fitch et al., 2015;Rabbani et al., 2016a). In low-level extent of modification of albumin by MG in experimental investigations *in vitro* and similar low-level extent of modification by MG of human serum albumin found similarly *in vivo*, MG was detected on these residues preferentially (Ahmed et al., 2005).

Applying RBD analysis and inspecting sequences for arginine or lysine residues at positions -4, -3, +3 and +4 with respect to the target arginine, we found the following number of arginine residues reactive towards MG modification and protein inactivation in predicted functional domains in SARS-CoV-2 proteins: nsp1, 2; nsp2, 3; nsp3, 3; nsp8, 1; nsp12, 1; nsp15, 2; spike protein, 2; M-protein, 1; NC, 10; and ORF10, 1. There were 25 functional arginines potentially activated for MG modification: 5 sites were in predicted α-helices and 2 in NC with predicted pK_a_ lowered by neighboring group interaction and thereby activated towards MG-modification (**Table 2**).

**Table 2.**
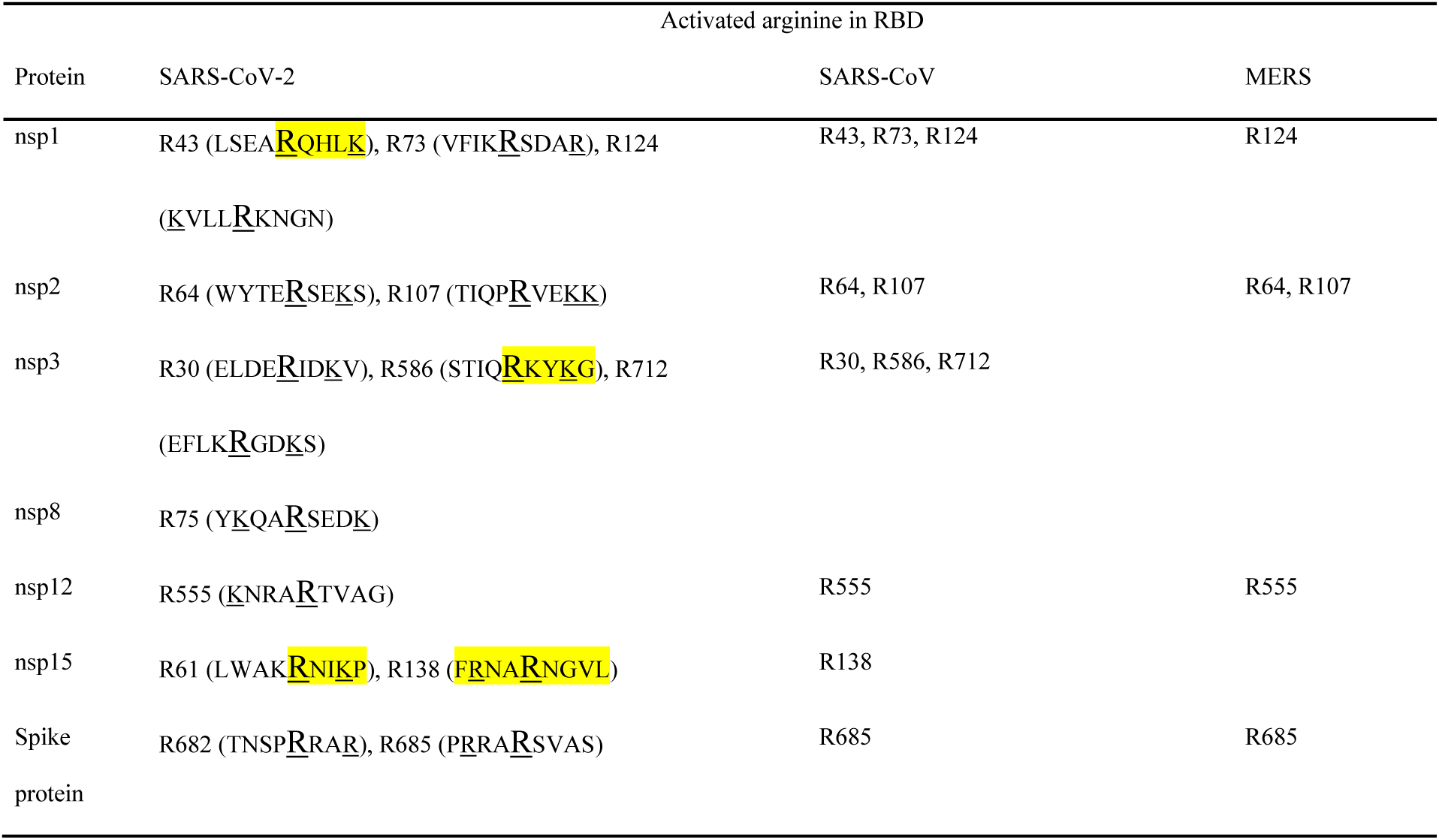
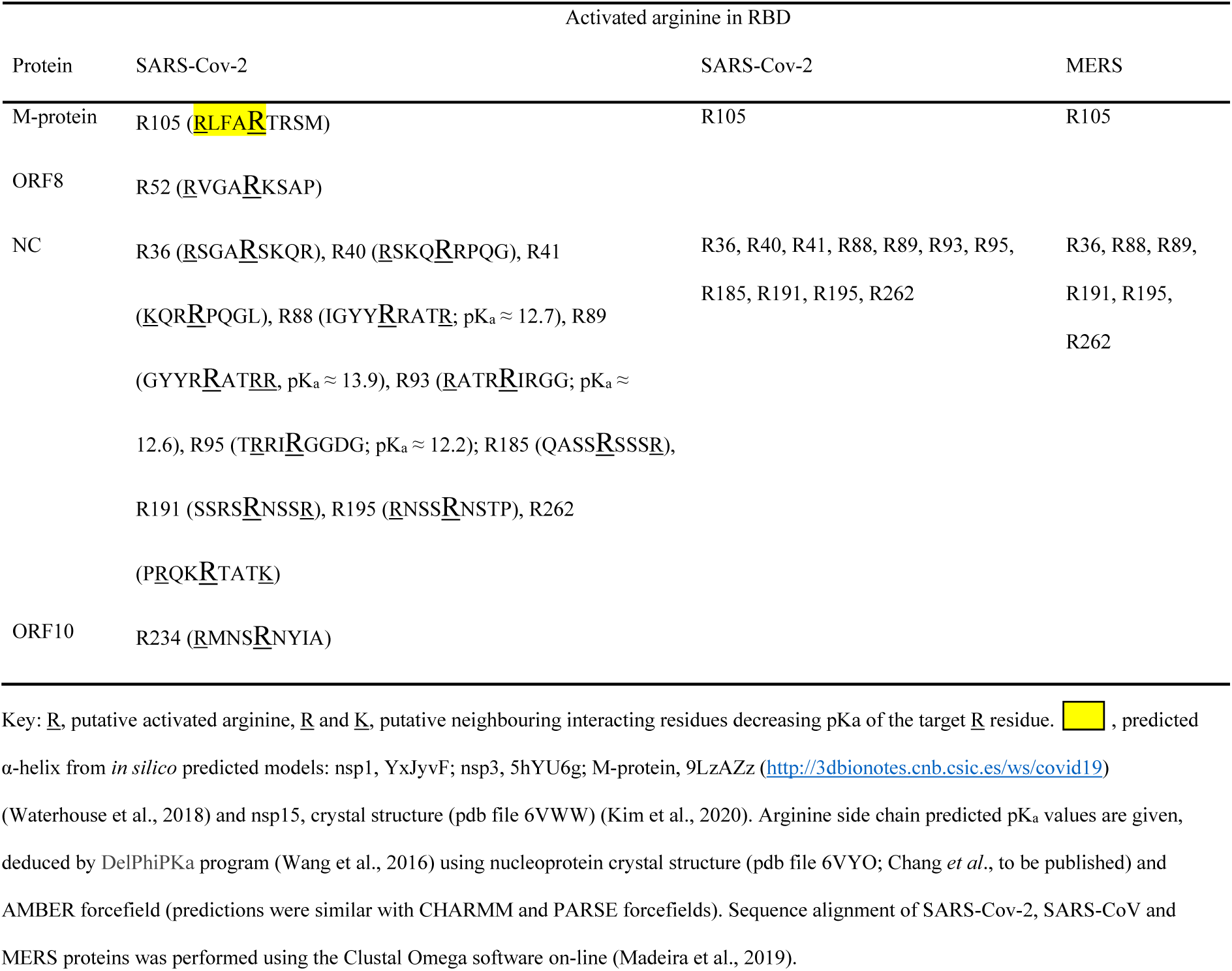
SARS-Cov-2, SARS-Cov and MERS proteins with putative activated arginine residues in functional domains.

Uniquely for related coronaviridae, SARS-CoV-2 spike protein has an interacting arginine triad, R_682_RAR_685_, at the S1/S2 cleavage site (Andersen et al., 2020) with both R682 and R685 predicted sites susceptible to MG modification (**Figure 2A**). Modification of this triad by MG is expected to confer resistance to proteolytic cleavage by transmembrane serine proteases (TMPRSSs) and blocking cell fusion for virion entry into pulmonary alveolar epithelial and other cell target sites, uncoating and replication (Meng et al., 2020). Trapped in the extracellular environment, there is excepted to be an improved host immune response to the virus; *cf*. viral host immunity boosted by similar aldehyde-modifying agents (Herrera-Rodriguez et al., 2019).

**Figure 2.**
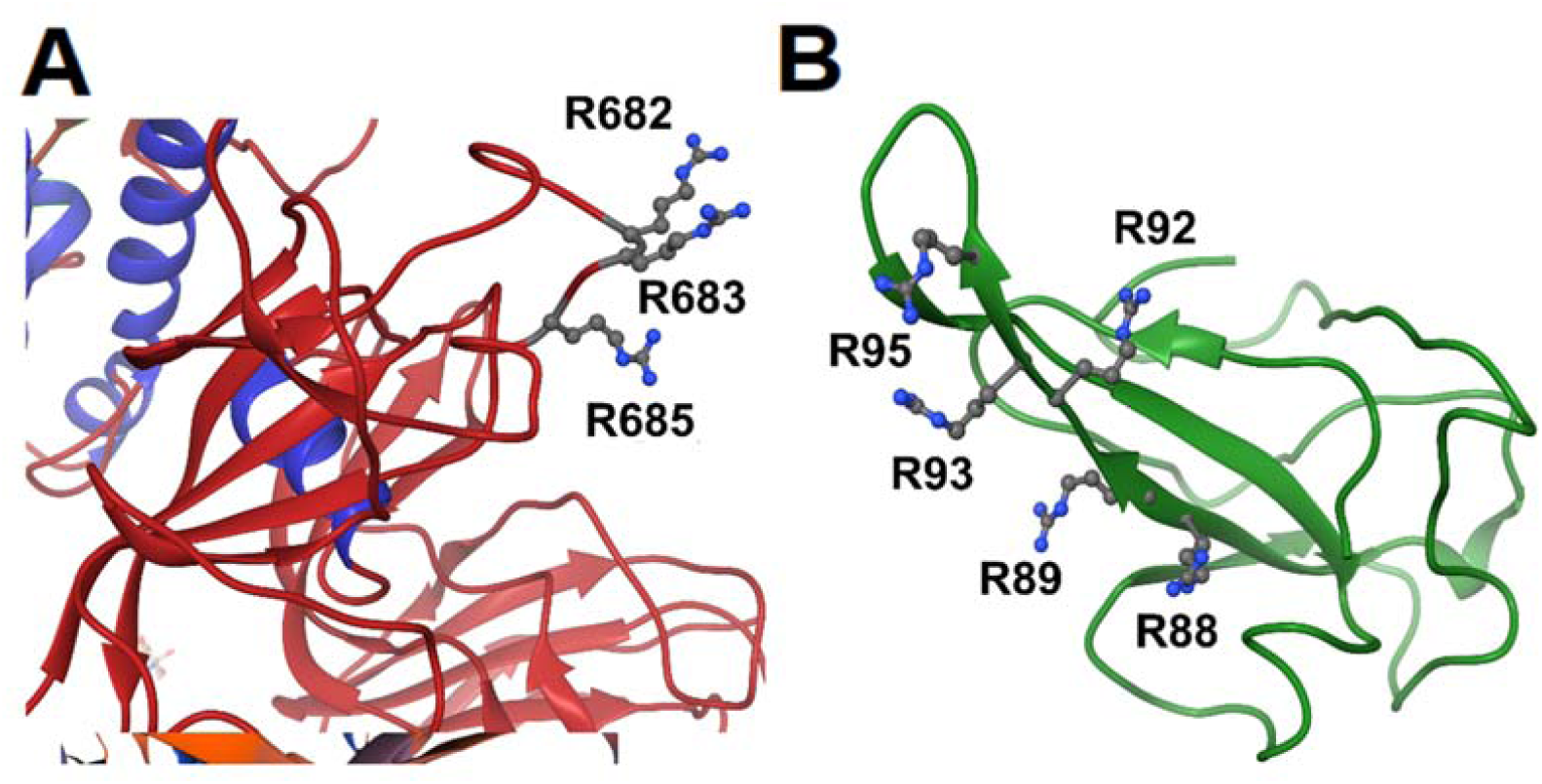
Activation of functional arginine residues towards modification by methylglyoxal in the SARS-CoV-2 proteome. **(A)** Activated arginine residues triad of Spike protein, R_682_R_683_AR_685_. (**B**) Activated arginine residue pentad of nucleoprotein, R_88_R_89_ATR_92_R_93_IR_95_. Molecular graphics produced from Spike protein (pdb file 6VSB) (Wrapp et al., 2020) and nucleoprotein segment crystal structure (pdb file 6VYO; Chang *et al*., to be published) using Chimera 1.14 (Pettersen et al., 2004).

SARS-CoV-2 nucleoprotein is highly susceptible to modification and functional inactivation by MG. Nucleoprotein binds the 3’ end of the viral single strand RNA genome of SARS-CoV-2 and it is arginine-rich, as is typical of RNA-binding proteins (Tan and Frankel, 1995). By analogy with SARS-CoV, residues 42-187 are involved with RNA binding with R93 playing a critical role (McBride et al., 2014). It is predicted to have sites susceptible to MG modification. The crystal structure of a segment of nucleoprotein, residues 50 – 173, is available and this enabled prediction of pK_a_ values of arginine residues in this region. pK_a_ shifts from 13.8 of inactivated arginine (Fitch et al., 2015) for R93 and R95 indicated *ca*. 16 and 40-fold increased reactivity towards MG modification, compared to non-activated target residue. These residues lie in a pentad of reactive arginine residues where MG modification at R93 and R95 is expected to inactive the nucleoprotein (**Figure 2B**). The SR-rich region of 182 – 196 is important for virus replication (Tylor et al., 2009) and is also a target for MG modification and inactivation at 3 sites: R185, R 191 and R195. MG modification of the nucleoprotein, and also membrane protein, will block viral replication and virion assembly, respectively.

There are similar MG modification sites in functional domains of related *coronaviridae*. The proteome of severe acute respiratory syndrome coronavirus (SARS-CoV) had 23 similar MG modification sites to those of SARS-CoV-2, and MERS coronavirus proteome had 12 similar MG modification sites. In all of these *coronaviridae* there were multiple MG modification sites in functional domains of the nucleoprotein (**Table 2**). Given the high activation of multiple arginine residues in functional domains of the SARS-CoV-2 proteome, it is likely that pharmacological increase of endogenous MG concentration will produce modification at multiple susceptible and functional sites, producing protein inactivation and antiviral response.

### Pharmacological increase of cellular methylglyoxal to virucidal levels by cell permeable glyoxalase 1 inhibitor and clinical antitumor drugs, doxorubicin and paclitaxel

Antiviral activity of supraphysiological concentrations of MG was reported historically (De Bock et al., 1957). More recently, inhibition of cytopathic effect of strains of influenza B by MG was investigated. The most sensitive strain gave a median inhibitory concentration of 23 ± 7 µM MG (Charyasriwong et al., 2016). These studies assessed antiviral activity by effect assessed by cytopathic response – concentration of MG required to prevent 50% lysis of infected cells, using relatively high multiplicity of infection (MOI). Median effective concentrations for antiviral effects of pharmacological agents tend to be lower in physiologically relevant range of MOI than in pathogenic response assessment (Wang et al., 2020b). These studies also used exogenous MG in cellular *in vitro* models where MG is rapidly metabolized by Glo1 and onward through the glyoxalase pathway to D-lactate (Rabbani et al., 2016b). The cellular concentration of MG is 1 – 4 µM and the plasma concentration 130 – 250 nM (Rabbani and Thornalley, 2014;Xue et al., 2016;Irshad et al., 2019). The optimum approach to achieve an antiviral effect is to increase cellular MG concentration by inhibition of Glo1. BBGD is a potent cell permeable Glo1 inhibitor prodrug. It delivers the Glo1 competitive inhibitor, S-p-bromobenzylglutathione (K_i_ = 160 nM), into cells and has established antitumor and antimalarial activity (Thornalley et al., 1994;Thornalley et al., 1996) (**Figure 3A**). We studied the effect of BBDG and clinically approved antitumor drugs on the cellular concentration of MG in human HEK293 cells – a cellular model employed for SARS-CoV-2 propagation (Chien et al., 2016). BBGD increased the endogenous concentration of cellular MG by 4-fold to *ca*. 20 µM – a level similar to that which inhibited viral cytopathic activity (Charyasriwong et al., 2016).

**Figure 3.**
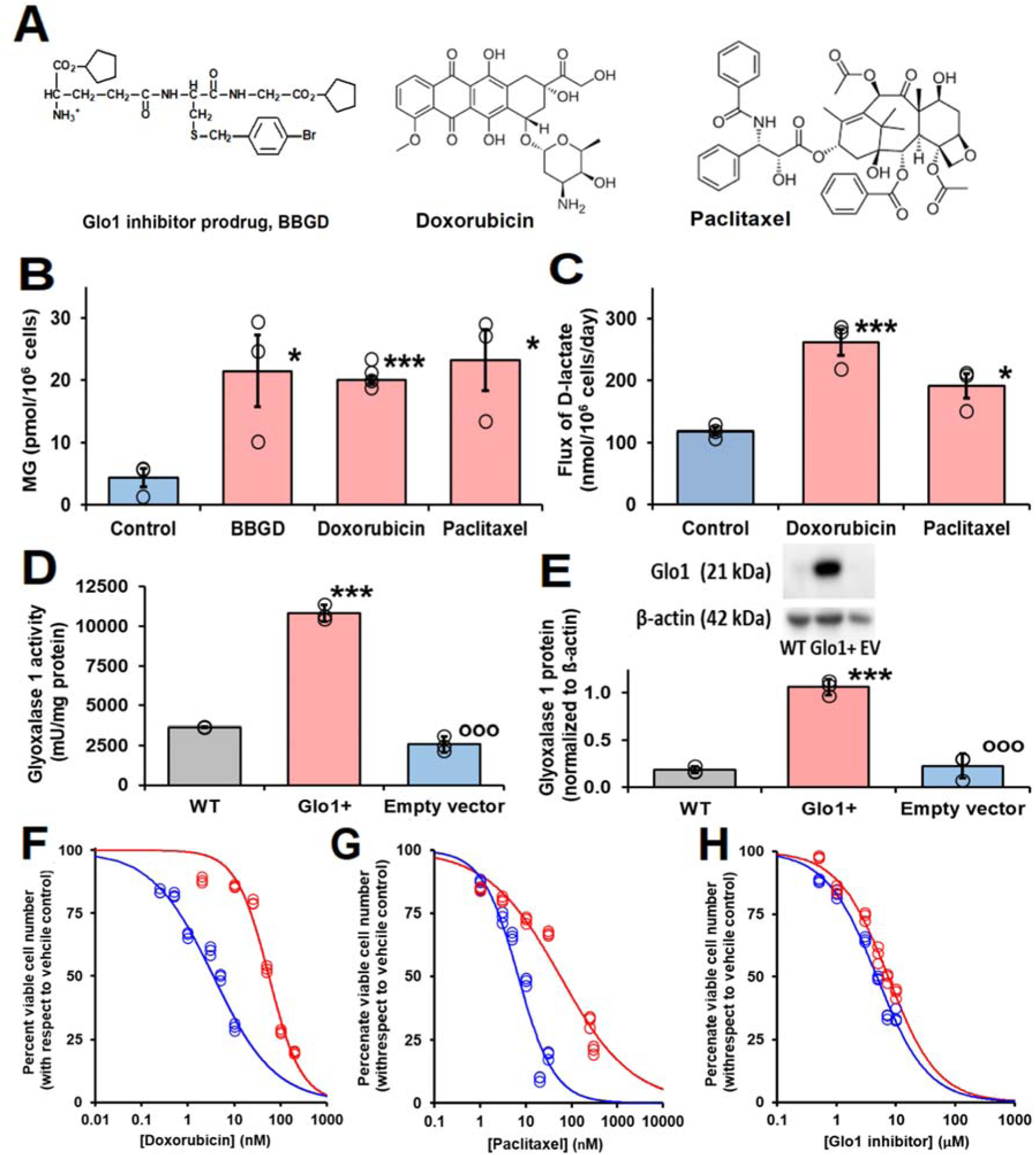
Glyoxalase 1 inhibitor prodrug, doxorubicin and paclitaxel increase cellular concentration of methylglyoxal to virucidal levels. **(A)** Molecular structures of drugs. Glyoxalase 1 inhibitor prodrug, S-p-bromobenzylglutathione cyclopentyl diester (BBGD). Delivers competitive Glo1 inhibitor, S-p-bromobenzylglutathione, K_i_ = 160 nM, into cells (Allen et al., 1993;Thornalley et al., 1996). Doxorubicin – topoisomerase inhibitor.(Tewey et al., 1984) Paclitaxel – stabilizer of microtubule assembly (Schiff et al., 1979). **(B)** and **(C)** Increase in cellular MG in HEK293 cells and flux of formation of D-lactate (surrogate for flux of MG), respectively, incubated *in vitro* with and without investigational agent and drugs indicated. For assay MG, cells were incubated with and without treatment for 3 h and for flux of D-lactate incubated for 24 h. Data are mean ± SEM (n = 3 except n = 4 for MG estimation with Doxorubicin; individual data points are shown). Drug concentrations: BBGD, 7.4 µM; doxorubicin, 6.0 nM, paclitaxel, 21 nM. Significance: b. P<0.02 and c. P<0.01 (*one-way ANOVA*) and * and ***, P<0.05 and P<0.001 with respect to control (*t-test*). (**D)** and (**E)** Activity and protein of Glo1, respectively, in HEK293 cells: wild-type (WT) and cells stably transfected to overexpress Glo1 (Glo1+) and empty vector (EV). Glo1 activity and protein were increased 4 – 5 fold. This was maintained for > 10 passages. **(F)** – **(H)**. Effect of Glo1 expression on anti-proliferative activity. Key: blue – empty vector, red – Glo1 overexpression. HEK293 cells were incubated with and without treatment for 48 h. Data (6 drug concentrations in triplicate) were fitted by non-linear regression to the dose-response equation, V = 100 x GC_50_^n^/(GC_50_^n^ + [Drug]^n^), solving for GC_50_ and, n (logistic regression coefficient). **(F)** Doxorubicin: empty vector, GC_50_ = 3.54 ± 0.28 nM, n = 0.71 ± 0.05; Glo1+, GC_50_ = 55.9 ± 3.4 nM, n = 1.24 ± 0.10 (16-fold resistance). **G** Paclitaxel: empty vector, GC_50_ = 6.8 ± 1.0 nM, n = 1.07 ± 0.17; and Glo1+, GC_50_ = 56.4 ± 7.2 nM, n = 0.55 ± 0.04 (8-fold resistance). **(H)** BBGD: GC_50_ = 4.78 ± 0.18 µM, n = 1.02 ± 0.05; and Glo1+, GC_50_ = 7.37 ± 0.30 µM, n = 1.04 ± 0.06 (2-fold resistance).

We have a longstanding interest in anticancer activity of Glo1 inhibitors and overexpression of Glo1 in multidrug resistant tumors (Thornalley et al., 1996;Rabbani et al., 2018). Hence, we have been studying the likely involvement of increased MG in the mechanism of action of clinical antitumor drugs. Interestingly, we found clinical antitumor agents, doxorubicin and paclitaxel (**Figure 3A**), also increased cellular MG by a similar extent (**Figure 3B**). Increased MG concentration induced by doxorubicin and paclitaxel is linked to increased glucose metabolism and related increased formation of MG as a byproduct of glycolysis. Indeed, flux of formation of D-lactate - a surrogate indicator of flux of formation of MG - was increased by both drugs (**Figure 3C**).

Increase of cellular MG also likely contributes to the antiproliferative effect of BBGD (Rabbani et al., 2018). The involvement of MG in the antiproliferative activity of doxorubicin and paclitaxel is unknown. We explored this by determining the effect of overexpression of Glo1 on inhibition of HEK293 cell growth. Vector-derived stable transfectant HEK293 cells lines were prepared with Glo1 expression increased 4 – 5 fold and empty vector transfectant control (**Figure 3D and 3E**), imposing a 4 – 5 fold increased rate of MG metabolism in Glo1 overexpressing cells. When these transfectant cell lines were treated with growth inhibitory concentrations of drugs, there was an increase of median growth inhibitory concentration GC_50_ value and resistance to inhibition of cell growth in HEK293 cells with stable overexpression of Glo1 (**Figure 3F and 3H**). The GC_50_ values were (mean ± SD; empty vector vs Glo1+): doxorubicin, 3.54 ± 0.28 nM vs 55.9 ± 3.4 nM (16-fold resistance); paclitaxel – 6.8 ± 1.0 nM vs 56.4 ± 7.2 nM (8-fold resistance). For treatment with BBGD there was an antiproliferative effect with limited change in GC_50_ value with Glo1 overexpression: 4.78 ± 0.18 µM vs 7.37 ± 0.30 µM (2-fold resistance). The limited effect on antiproliferative effect of BBGD is expected as the delivered Glo1 inhibitor also inhibits the overexpression factor, Glo1.

## DISCUSSION

The enrichment of arginine residues in functional domains of the SARS-CoV-2 proteome provides important evidence to support an arginine-modifying agent strategy for inactivation of the virus and virucidal activity. Arginine residues are also enriched in the human host proteome but less so that in SARS-CoV-2; 3.6 versus 4.9. This characteristic of the human proteome was noted previously (Gallet et al., 2000) – now updated herein with UniProtKB current sequence information. In addition, in the SARS-CoV-2 proteome there is a high number of arginine residues activated by neighbouring groups for reaction with MG. This particularly applies to the nucleoprotein and, uniquely for the SARS-CoV-2 coronavirus, to the furin cleavage site of the spike protein. For the 25 arginine residues identified with predicted reactivity towards MG modification in the SARS-CoV-2 proteome, there is supporting secondary structure and predicted low pK_a_ value evidence for 7 of them. These arginine residue targets are in key proteins: nucleoprotein, M-protein and Spike protein. A further important feature for susceptibility of viral proteins to MG modification is protein abundance: high abundance of a protein increases its susceptibility to reaction with MG. Previous earlier studies of the SARS virion suggested proteins of highest abundance were: nucleoprotein, M-protein, Spike protein and nsp3 (Neuman et al., 2008). Assuming a similar relative abundance of proteins in SARS-CoV-2, multiple arginine residues reactive towards modification and inactivation by MG were found in the 4 most abundant proteins of the SARS-CoV-2 proteome.

We also explored use of proteomics data from previous studies where MG modification was detected at 411 in different sites in the cytosolic extract of human endothelial cells in culture (Irshad et al., 2019) in an attempt to identify a proteomic MG modification motif to map onto the SARS-CoV-2 proteome (data not shown). Although we could determine amino acid frequencies round the MG-modified arginine targets, there was a dropout of peptides (failure to detect) without lysine and arginine residues on the N-terminal side of the arginine target in mass spectrometric detection. In comparisons with unmodified arginine target peptide sequences, this led to an artifactual enrichment of lysine and arginine residues on the N-terminal side of the MG-modified arginine target. This peptide dropout was likely due to an additional missed cleavage by trypsin when the target arginines were modified by MG, making the related tryptic peptides difficult to detect in mass spectrometry analysis due to the loss of arginine residue charge and increased sequence length. This requires further investigation and likely studies with other proteases other than trypsin in which MG modification changes tryptic peptide formation.

SARS-CoV-2 was rich with arginine residues in functional sites activated to MG modification. Other *coronaviridae -* SARS-CoV-2 and MERS - had similar MG modification sites in protein crucial for virion viability – particularly the nucleoprotein. This suggests that pharmacological agents increasing cellular concentration of MG, inducing dicarbonyl stress, may have virucidal activity against multiple *coronaviridae*.

Doxorubicin and paclitaxel are clinical antiproliferative antitumor agents with mechanisms of action targeting inhibition of topoisomerase-II in DNA replication and stabilization of the interphase and microtubular network and mitotic spindle in mitosis, respectively (Schiff et al., 1979;Tewey et al., 1984). Herein, we show that increase in MG contributes to their mechanism of antiproliferative activity. Doxorubicin increases glucose metabolism by increasing expression of glucose transporter GLUT1 and hexokinase-2 (Demel et al., 2015). Paclitaxel stabilizes microtubules, decreasing free tubulin concentration; the latter increasing mitochondrial voltage-dependent anion channel (VDAC) activity and thereby *in situ* activity of hexokinase (Maldonado et al., 2010). These mechanisms are available in the lung epithelial cells primarily targeted by SARS-Cov-2 (Pezzulo et al., 2011;Lottes et al., 2014). Increased glucose metabolism produces a corresponding increase in the formation of MG – evidenced herein by increase in flux of formation of D-lactate; there may be disproportionately large increase in MG if expression of enzymes of onward metabolism of triosephosphates, triosephosphate isomerase and glyceraldehyde-3-phosphate dehydrogenase, are not increased along with hexokinase activity and glycolysis becomes dysregulated or unscheduled (Irshad et al., 2019;Rabbani and Thornalley, 2019).

In the search for drugs to repurpose for COVID-19 disease, we suggest doxorubicin and paclitaxel be considered. These drugs have not been proposed hitherto although they have been evaluated for antiviral activity, particularly with respect to inhibition of viral helicase (Ash and Diekema, 1987;Bergamini et al., 1992;Borowski et al., 2002;Briguglio et al., 2011). Paclitaxel also suppressed inflammation in a murine model of bacterial pneumonia (Mirzapoiazova et al., 2007). However, a concern is the established adverse effects of these drugs found in cancer chemotherapy: bone marrow suppression (primarily neutropenia) and peripheral neuropathy for paclitaxel, and cumulative congestive heart failure for doxorubicin. Toxicity is related to dose and duration of treatment (Rowinsky, 1997;Barrett-Lee et al., 2009). Drug treatment of COVID-19 may be shorter than in cancer chemotherapy: for example, median hospitalization time of patients surviving severe symptoms of COVID-19 was 28 days (Wang et al., 2020a) and a typical course of cancer chemotherapy with paclitaxel and doxorubicin is 6 months or longer (Rowinsky, 1997;Barrett-Lee et al., 2009). If high potency antiviral effect of these agents is found, low dose and short duration of treatment is expected to decrease risk of adverse effects.

The approach to drug repurposing for COVID-19 developed herein addresses the intrinsic vulnerability of SARS CoV-2 proteome to endogenous reactive metabolites, with respect to the human host, and identified drugs to exploit this. Other strategies for repurposing drugs are based on SARS CoV-2 protein interactions with human host proteins and drugs targeted to them, virion endosomal processing and viral protease inhibition (Chen et al., 2020;Gordon et al., 2020;Wang et al., 2020b).

## CONCLUSIONS

We provide evidence of vulnerability of SARS-CoV-2 to modification and inactivation by MG. We also reveal, for the first time, increase in cellular concentration of MG in the antiproliferative activity of doxorubicin and paclitaxel – thereby providing a mechanistic rationale for repurposing of these drugs against SARS-CoV-2 and treatment of COVID-19 disease. Doxorubicin and paclitaxel may have potential for application for treatment of COVID-19 and may now be considered for evaluation in SARS-CoV-2 live virus cultures and animal models.

## DATA AVAILABILITY STATEMENT

All datasets presented in this study are included in the article and Supplementary Material.

## ETHICS STATEMENT

All methods were carried out in accordance with relevant guidelines and regulations and all experimental protocols were approved by University of Warwick Genetic Modification & Biosafety Committee (Project no. 305).

## AUTHOR CONTRIBUTIONS

MA-M accessed protein sequence information on the SARS-CoV-2, applied RBD analysis and produced the molecular graphics images. HA cultured HEK293 cells, prepared and propagated plasmids, prepared stable transfectant cell lines and performed metabolite and drug treatment studies. PW and AdlF collated and curated data on arginine sequence environments. MX provided technical guidance and support to HA and performed SARS-CoV2, SARS-CoV and MERS sequence alignments. PJT assisted with MG analysis. NR and PJT acquired the funding, designed and supervised the studies, contributed to the data analysis and wrote the manuscript. All authors read, edited and approved the final manuscript.

## SUPPLEMENTARY MATERIAL

The Supplementary Material for this article can be found online.

## SUPPLEMENTARY TABLES

**Table S1.** SARS-CoV-2 viral proteome.

| No | Protein (acronym(s)) | Reference sequence<br>(no of a.a. residues) | Function |
| --- | --- | --- | --- |
| 1. | Non-structural protein 1<br>(nsp1) | YP_009725297.1<br>(180 residues) | Inhibits host translation by interacting with the 40S ribosomal subunit. |
| 2. | Non-structural protein 2<br>(nsp2) | YP_009725298.1<br>(638 residues) | May modulate host cell survival signaling pathway by interacting with host prohibitin-1 and prohibitin-2. |
| 3. | Non-structural protein 3<br>/Papain-like proteinase<br>(nsp3, PL-PRO) | YP_009725299.1<br>(1945 residues) | Cleaves N-terminus of the replicase polyprotein. With nsp4, participates in assembly of viral membrane assembly. Antagonizes type I interferon innate immunity by. Prevents host NF-kappa-B signaling. |
| 4. | Non-structural protein 4<br>(nsp4) | YP_009725300.1<br>(500 residues) | Participates in the virion double-membrane vesicles assembly. |
| 5. | Non-structural protein 4<br>(nsp5, 3C-like proteinase,<br>3CL-PRO, Main protease) | YP_009725301.1<br>(306 residues) | Cleaves C-terminus of replicase polyprotein R1ab at 11 sites. |
| 6. | Non-structural protein 6<br>(nsp6) | YP_009725302.1<br>(290 residues) | Role in initial induction and regulation of autophagosomes from host reticulum endoplasmic. |
| 7. | Non-structural protein 7<br>(nsp7) | YP_009725303.1<br>(83 residues) | Forms a hexadecamer with nsp8 that may participate in viral replication by acting as a primase or may synthesize longer oligonucleotides. |
| 8. | Non-structural protein 8<br>(nsp8) | YP_009725304.1<br>(198 residues) | Forms a complex with nsp7 that may act as a primase or may synthesize longer oligonucleotides. |
| 9. | Non-structural protein 9<br>(nsp9) | YP_009725305.1<br>(113 residues) | May be an ssRNA-binding protein participating in viral replication. |
| 10. | Non-structural protein 10<br>(nsp10) | YP_009725306.1<br>(139 residues) | Pivotal role in viral transcription by stimulating both nsp14 3'-5' exoribonuclease and nsp16 2'-O-methyltransferase activities for viral mRNAs cap methylation. |
| 11. | Non-structural protein 11<br>(nsp11) | YP_009725312.1<br>(13 residues) |  |

**Table S1 | SARS-CoV-2 viral proteome (cont'd).**
| No | Protein (acronym(s)) | Reference sequence<br>(no of a.a. residues) | Function |
| --- | --- | --- | --- |
| 12. | RNA-directed RNA polymerase (nsp12, Pol/RdRp) | YP_009725307.1<br>(932 residues) | Replication and transcription of the viral RNA genome. Part of orf1ab polyprotein. |
| 13. | Helicase (nsp13, Hel) | YP_009725308.1<br>(601 residues) | Multi-functional protein with RNA and DNA duplex-unwinding activities with 5' to 3' polarity. |
| 14. | Guanine-N7 methyl-transferase (nsp14, ExoN) | YP_009725309.1<br>(527 residues) | Exoribonuclease activity acting on both ssRNA and dsRNA in a 3' to 5' direction. N<sup>7</sup>-guanine methyltransferase activity. |
| 15. | Uridylate-specific endoribonuclease (nsp15, NendoU) | YP_009725310.1<br>(346 residues) | Mn<sup>2+</sup>-dependent, uridylate-specific endoribonuclease. |
| 16. | 2'-O-ribose methyl-transferase (nsp16) | YP_009725311.1<br>(298 residues) | Methyltransferase mediating mRNA cap 2'-O-ribose methylation to the 5'-cap structure of viral mRNAs - prerequisite for binding of nsp16. Essential function to evade immune system. |
| 17. | Spike glycoprotein (Spike protein S1, S-glycoprotein) | YP_009724390.1<br>(1273 residues) | Attaches the virion to the cell membrane by interacting with host receptor, initiating the infection. Binding to human ACE2 and CLEC4M/DC-SIGNR receptors. |
| 18. | Protein 3a (ORF3a) | YP_009724391.1<br>(275 residues) | Forms K<sup>+</sup>-sensitive ion channels which may modulate virus release. Up-regulates fibrinogen subunit expression in host lung epithelial cells. Downregulates the type 1 interferon receptor. |
| 19. | Protein 3b (ORF3b)* | (57 residues) | Inhibits expression of β-interferon. |
| 20. | Envelope small membrane protein (E protein) | YP_009724392.1<br>(75 residues) | Viroporin in host membranes forming pentameric protein-lipid pores allowing ion transport. Also involved in the induction of apoptosis. |
| 21. | Membrane protein (M protein) | YP_009724393.1<br>(222 residues) | Component of the viral envelope involved in virus morphogenesis and assembly. |
| 22. | Non-structural protein 6 (ns6, ORF6) | YP_009724394.1<br>(61 residues) | Determinant of virus virulence. |

**Table S1 | SARS-CoV-2 viral proteome (cont'd).**
| No | Protein (acronym(s)) | Reference sequence<br>(no of a.a. residues) | Function |
| --- | --- | --- | --- |
| 23. | Protein 7a (ORF7a) | YP_009724395.1<br>(121 residues) | Non-structural protein of unknown function |
| 24. | Protein non-structural 7b<br>(ORF7b) | YP_009725318.1<br>(43 residues) | Unknown function |
| 25. | Protein non-structural 8<br>(ORF8) | YP_009724396.1<br>(121 residues) | May play a role in host virus interaction. |
| 26. | Nucleoprotein (NC, ORF9) | YP_009724397.2<br>(419 residues) | Packages the positive strand viral genome RNA into a helical ribonucleocapsid. Also enhances efficiency of viral RNA transcription and viral replication. |
| 27. | Protein 9b (ORF9b) | ORF9B_WCPV<br>(97 residues) | Unknown function. |
| 28. | ORF10 protein | YP_009725255.1<br>(38 residues) | Unknown function. |
| 29. | Uncharacterized protein 14<br>(ORF14) | Y14_WCPV<br>(73 residues) | Unknown function. |

\*Sequence from reference (Chan et al., 2020).

**Table S2.** Receptor binding domain of SARS-CoV-2 viral proteome.

| Amino acid | N |  | Prevalence |  | Proportion | Fold |
| --- | --- | --- | --- | --- | --- | --- |
|  | All | (RBD) | All | RBD | in RBD (%) | Enrichment |
| Ala | 667 | 18 | 6.76 | 2.9 | 2.7 | 0.4 |
| Arg | 358 | 110 | 3.63 | 17.7 | 30.7 | 4.9 |
| Asn | 537 | 42 | 5.44 | 6.8 | 7.8 | 1.2 |
| Asp | 514 | 45 | 5.21 | 7.3 | 8.8 | 1.4 |
| Cys | 310 | 15 | 3.14 | 2.4 | 4.8 | 0.8 |
| Gln | 369 | 29 | 3.74 | 4.7 | 7.9 | 1.3 |
| Glu | 452 | 41 | 4.58 | 6.6 | 9.1 | 1.4 |
| Gly | 574 | 22 | 5.82 | 3.5 | 3.8 | 0.6 |
| His | 193 | 17 | 1.96 | 2.7 | 8.8 | 1.4 |
| Ile | 514 | 8 | 5.21 | 1.3 | 1.6 | 0.3 |
| Leu | 950 | 27 | 9.63 | 4.4 | 2.8 | 0.5 |
| Lys | 568 | 74 | 5.76 | 11.9 | 13.0 | 2.1 |
| Met | 203 | 7 | 2.06 | 1.1 | 3.5 | 0.6 |
| Phe | 504 | 13 | 5.11 | 2.1 | 2.6 | 0.4 |
| Pro | 404 | 23 | 4.09 | 3.7 | 5.7 | 0.9 |
| Ser | 665 | 53 | 6.74 | 8.5 | 8.0 | 1.3 |
| Thr | 728 | 33 | 7.38 | 5.3 | 4.5 | 0.7 |
| Trp | 113 | 2 | 1.15 | 0.3 | 1.8 | 0.3 |
| Tyr | 448 | 23 | 4.54 | 3.7 | 5.1 | 0.8 |
| Val | 796 | 18 | 8.07 | 2.9 | 2.3 | 0.4 |
| Total: | 9867 | 620 | 100 | 100 |  |  |

